# Rational engineering of low temperature activity in thermoalkalophilic *Geobacillus thermocatenulatus* lipase

**DOI:** 10.1101/2021.03.14.435354

**Authors:** Weigao Wang, Siva Dasetty, Sapna Sarupria, Mark Blenner

**Affiliations:** Department of Chemical & Biomolecular Engineering, Clemson University, Clemson, SC 29634; Department of Chemistry, University of Minnesota, Minneapolis, MN, 55455; Department of Chemical & Biomolecular Engineering, 590 Avenue 1743, University of Delaware, Newark, DE 19713

**Keywords:** robust enzyme, temperature adaptation, cold activity, active site flexibility, *Geobacillus thermocatenulatus*, lipase

## Abstract

While thermophilic enzymes have thermostability desired for broad industrial applications, they can lose activity at ambient temperatures far from their optimal. Engineering cold activity into thermophilic enzymes has the potential to broaden the range of temperatures resulting in significant activity (i.e., decreasing the temperature dependence of k_cat_). Even though it has been widely suggested that cold temperature enzyme activity results from active flexibility that is at odds with the rigidity necessary for thermostable enzymes; however, directed evolution experiments have shown us these properties are not mutually exclusive. In this study, rational protein engineering was used to introduce flexibility inducing mutations around the active sites of *Geobacillus thermocatenulatus* lipase (GTL). Two mutants were found to have enhanced specific activity compared to wild-type at temperatures between 283 K to 363 K with p-nitrophenol butyrate but not with larger substrates. Kinetics assay revealed both mutations resulted in psychrophilic traits, such as lower activation enthalpy and more negative entropy values compared to wild type in all substrates. Furthermore, the mutants had significantly improved thermostability compared to wild type enzyme, which proves that it is feasible to improve the cold activity without trade-off. Our study provides insight into the enzyme cold adaptation mechanism and design principles for engineering cold activity into thermostable enzymes.

## 1.0 Introduction

Extremophile enzymes are of central interest to understand the molecular origin of the relation between enzyme activity-stability and temperature [1–6]. Their unique properties have been exploited across the food, chemical and pharmaceutical industries [7]. Thermophilic enzymes are usually found in microorganisms growing at high temperature (between 45°C and 121 °C). To retain functional activity, thermophilic enzymes have evolved structural traits such as charged surface residues, large hydrophobic cores, shortened loops and disulfide bonds, which provide larger barriers to thermally induced unfolding [8, 9]. In contrast, psychrophilic enzymes have no such evolutionary pressures on stability [10, 11]. They do, however, have to defy the exponential activity-temperature relationship to function at low temperatures (< ~20°C) with rates similar to meso- and thermophiles at their optimal temperatures [12, 13]. Structural features contributing to the cold activity include a smaller number of salt bridges, a reduced number of disulfide bonds and increased frequency of small and neutral amino residues [2, 14–16].

The relationship between the psychrophilic enzyme structural traits and cold adaptation have been discussed systematically before [2, 12, 17–22]. According to Feller et al., psychrophilic enzymes tend to have lower activation enthalpy (ΔH^‡^) and more negative activation entropy (Δ*S*^‡^) values [19]. The lower activation enthalpy and entropy are believed to be the primary adaptations in psychrophilic enzymes that make the reaction rate less temperature-dependent [23]. Reflected on the structure, psychrophilic enzymes are believed to compensate for increased flexibility near the active site through rigidification in other parts of the protein [2, 19]. The increased flexibility in the reactant-bound-enzyme state would imply a larger configurational space and result in a more negative activation entropy (i.e., a greater loss of configurational entropy) if the transition state remains approximately the same as in the wild-type enzyme. This was supported by the higher measured Michaelis-Menten (*K_m_*) constants of several psychrophilic enzymes compared to their mesophilic counterparts, which is an indication of weak interactions between enzyme and substrate [19].

The notion that cold activity and thermostability in enzymes has been repeated throughout the literature [24]; however, Differential Scanning Calorimetry (DSC) studies of multi-domain protein phosphoglycerate kinase (PGK) showed that PGK a flexible thermolabile domain and a thermostable domain unfolded independently and effectively partitioned flexibility and thermostability [25]. It was then proposed that the thermostability and cold activity can evolve separately [17, 18, 25]. Therefore, engineering the active sites by increased flexibility could be a promising strategy to rationally change the temperature-dependence of the enzyme and make robust thermostable enzymes that can maintain activity across a wide range of temperatures [18]. Earlier studies of cold activity engineering in subtilisin-like protease relies on random mutagenesis and recombination [26]. One of the recent cold adaptation engineering via rational design was achieved on *Bacillus agaradherans* cellulase Cel5A [27]. In this study, activation energy, entropy and enthalpy values were reduced by disrupting the hydrogen bonds around the catalytic residues. The increased activity at low and warmer temperatures from the designed mutants were attributed to the increased fluctuation proximal to the active site [27].

In our study, *Geobacillus thermocatenulatus* lipase was used as a model enzyme to rationally engineer cold activity. We hypothesized that replacing residues around the active sites with smaller size amino acids could introduce flexibility to the active site. Most mutants showed insoluble expression and significant reduction in their specific activities. Of the designed mutants, only E316G and E361G were soluble and both exhibited improved specific activity than wild-type (WT) on substrate p-nitrophenol butyrate (C4). Kinetics experiment indicated that both mutants showed reduced activation enthalpy and entropy compared with WT, a cold-adaptation feature found in psychrophilic enzymes. Although the specific activity of mutants for longer substrates p-nitrophenol octanoate (C8) and p-nitrophenol laurate (C12) was less than the specific acitvity of the wild-type lipase, similar decrease in activation enthalpy and entropy was observed.

## 2.0 Materials and Methods

All chemicals used in this study were obtained from Sigma Aldrich. Cloning related enzymes were purchased from New England Biolabs (NEB). Zyppy^™^ Plasmid Miniprep kit and DNA Clean & Concentrator^™^ kit were purchased from Zymo Research. Gene fragment of GTL was codon optimized using COOL (http:///cool.syncti.org) and synthesized from Eurofins company. Oligos used for the site directed mutations were purchased from IDT DNA.

### 2.1 Cloning, mutagenesis and purification experiments of GTL

Plasmid propagation and subcloning was performed using Escherichia coli DH10β competent cells (NEB). The codon optimized GTL gBlock (Supporting Information Table 1) was cloned into pET15b vector using sequence and ligation independent cloning [1]. Transformation in chemically competent *E.coli* was performed using standard heat shock methods. Two step site directed mutagenesis (using primers described in Supporting Information Table 2) was used to introduce mutations [2]. In brief, PCR was performed with reaction mixture containing forward and reverse primer separately for 5 cycles and were combined together and for another 16 cycles using the same PCR protocol. All mutations were verified by Sanger sequencing. WT and mutant plasmids were miniprepped and transformed into *E. coli* BL21(DE3) competent cells (NEB). A single colony was picked and inoculated into 5 mL of LB+Amp medium in a 15mL falcon tube. Overnight cultures were then transferred into 1 L of LB+Amp medium in a 2.8 L baffled shake flask and incubated at 310 K, 225 rpm. When the OD_600_ reached 0.4-0.6, expression was induced at 291 K with 0.2 mM IPTG (isopropyl β-D-1-thiogalactopyranoside) for 10 hours. The cell pellets harvested and then sonicated in 50 mM Tris-HCl at pH 7.4 and 500 mM NaCl buffer and then clarified by ultracentrifugation. The filtered supernatant was loaded on to a Ni-nitrolotriacetate (Ni-NTA) column (GE, USA). The proteins were eluted with a 40mM imidazole buffer and buffer exchanged with 20 mM Tris-HCl buffer at pH 8.0 using 10 kDa cutoff Amicon ultrafilter (Millipore, USA). The final concentration (C) of purified was measured using NanoDrop A280 (Thermo Fisher, USA) according to Beer’s law: C = A * (MW)/ε_molar_, where A is the absorbance, MW is the molecular weight of the protein, the concentration is in mg/mL. The extinction coefficient ε_molar_ was calculated from the website (web.expasy.org).

### 2.2 Enzymatic activity assay

The lipase activity was measured using p-nitrophenol esters (p-nitrophenol butyrate, p-nitrophenol octanoate and p-nitrophenol laurate). Upon hydrolysis, the released p-nitrophenol (pNP) exhibits a characteristic maximum absorbance at 405 nm. A 1 ml reaction mixture was prepared for the enzymatic assay as follows: 950 μl of 50 mM Tris-HCl buffer at pH 8.0 containing pure enzyme, 40 μl pure ethanol, and 10 μl of 10 mM p-nitrophenol ester in acetonitrile. The blank mixture was prepared with enzyme-free solution. The reaction mixture containing substrate was incubated in mini dry bath (Fisher Scientific, USA) till it reached to the desired temperature (confirmed with a temperature monitored blank reaction), and then enzyme was added to trigger the reaction. Each reaction was incubated at a specified temperature in a mini heating block for 5 minutes, before rapidly moving to the plate reader for measurement. The released pNP (extinction coefficient ε=16,000 M^-1^cm^-1^) was measured at 405 nm using Epoch 2 plate reader (Biotek, US) set to 37°C and subtracting the blank reading. One unit of enzyme activity was defined as 1 μmol pNP liberated per minute per microgram enzyme. All measurements were made in at least triplicate.

### 2.3 Analysis of biochemical properties

Thermal stability was measured by incubating purified enzymes in a mini dry block at 343 K for 0 or 3 hours. The residual activity was determined using the standard enzymatic activity assay at 323 K. The fractional activities retained for both WT and mutants were calculated by dividing the 3 hour-incubation activities by the 0 hours-incubation activity. All measurements were made in at least triplicate.

The kinetics parameters (*V_max_, K_m_*) of the WT and mutants were determined by using Epoch 2 plate reader (BioTek, USA) at different temperatures ranging from 303 K to 338 K with 5 K increments. The absorbance values were corrected with extinction coefficients that are corrected for temperature dependence. Initial rates were determined using 0.1 μM enzyme in 200 μl buffer, and a range of substrate concentrations: from 50 μM to 800 μM for C4, and from 5 μM to 80 μM for C8 and C12. All measurements were made in at least triplicate. Kinetic parameters were determined by a non-linear least squares regression performed in Prism 6. The kinetic rate constant, *k_cat_*, was determined by dividing *V_max_* by the specific activity of each enzyme preparation. Van’t hoff plots were created by linear regression of a semi-log plot of *k_cat_* as a function temperature, taken over the linear range of data.

### 2.4 CD spectroscopy

CD measurements were performed with Jasco-J-810 CD spectropolarimeter (Jasco, Japan). The measured enzymes were dissolved in 10 mM potassium phosphate pH 7.0. 300 μL pure protein solution was added to a 1 mm pathlength quartz cuvette (Starna Cells, USA). The scanning wavelength starts from 300 nm and ends at 190 nm, with a 2nm decrement, accumulating 6 scans. Spectra were corrected for background by subtracting buffer-only spectra. The circular dichroism in units of milidegrees (mdeg) were converted into molar ellipticity using the equation: Θ= m°• 100/(L•C). m° is the raw output in mdeg, C is the protein concentration in mM, L is the cell pathlength in cm. The secondary structure content was calculated by spectral deconvolution using CDpro continll [3].

## 3.0 Results

### 3.1 Mutation Design

GTL is a large thermoalkalophilic lipase with a molecular weight of ~43 kDa and optimal activity at 338 K and pH 8-10 [28]. The three-dimensional structure of GTL belongs to the *α/β* hydrolase (Fig. 1a). It contains two cofactor ions (Zn^2+^ and Ca^2+^) and an amphipathic helix-loop motif that act as a lid protecting the active site. The catalytic triad is comprised of Ser114-His359-Asp318. Potential mutation sites were selected by inspection (PDB: 2W22) of the four residues on either side of the catalytic residue, ignoring residues that were located within non-loop secondary structure. The rationale was flexibility inducing mutations in secondary structures might cause a significant perturbation in the structure. This narrowed the potential residues for mutation to Glu316, Asn317, Trp314, His113, Ala112, Val357 and Glu361 (Fig. 1b). All these residues were mutated to Gly, and His113 and Glu361 were replaced with Ala and Asp separately. Mutations were designed based on the rule that it decreased the number of hydrogen bonds number (Table 1). Mutants E316G and E361G were selected due to their soluble expression.

**Figure 1.**
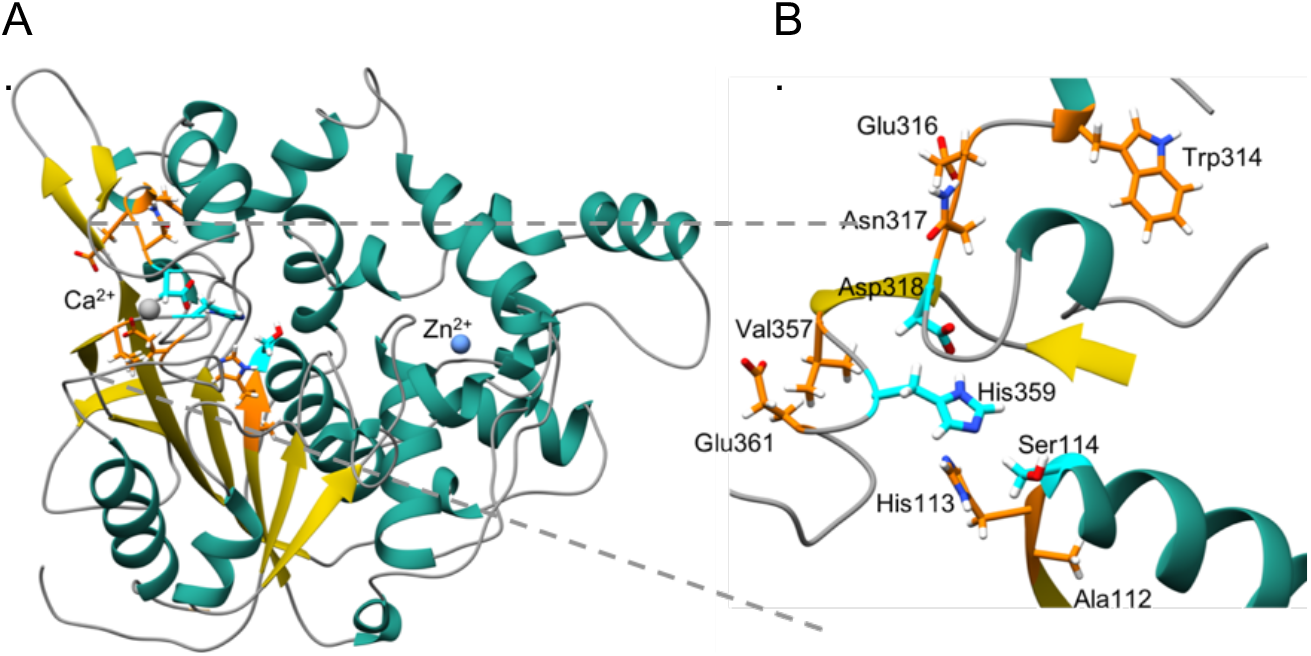
(a) Cartoon representation of the crystal structure of GTL (PDB: 2W22) with *α*-helices and *β*-sheets shown in teal and yellow, respectively. The active site residues are represented as ball and stick models, and its carbons are colored in cyan. Carbon atoms of sites for mutations selected to enhance active site flexibility are colored in orange, and represented by ball and stick model. Nitrogen, oxygen, and hydrogens are colored in blue, red and white, respectively. (b) Closer view of the region around active site of GTL.

**Table 1.** Hydrogen Bonds Before and After Mutations

|  |  |  |
| --- | --- | --- |
| A112G | Before | Ala112-Thr160, Ala112-Ile162 |
|  | After | None |
| H113A | Before | His113-His359, His113-Tyr30, His113-Leu14, His113-Gln115, His113-Gly116 |
|  | After | Ala113-Leu14, Ala113-Gln115, Ala113-Gly116 |
| H113G | Before | His113-His359, His113-Tyr30, His113-Leu14, His113-Gln115, His113-Gly116 |
|  | After | None |
| W314G | Before | Trp314-Asn322, Trp314-Asp311 |
|  | After | None |
| E316G | Before | Glu316-Thr323, Glu316-Tyr282 |
|  | After | None |
| N317G | Before | Asn317-Gly319 |
|  | After | None |
| N356G | Before | Asn356-Glu271, Asn356-Glu285 |
|  | After | None |
| V357G | Before | Val357-Glu271 |
|  | After | None |
| E361D | Before | Glu361-Gly364, Glu361-Asn368, Glu361-Asp358, Glu361-Arg272, Asp358-Asn289 |
|  | After | Asp361-G364, Asp358-Asp361 |
| E361G | Before | Glu361-Gly364, Glu361-Asn368, Glu361-Asp358, Glu361-Arg272, Asp358-Asn289 |
|  | After | None |

Note: “-” represents the hydrogen bond between residues.

### 3.2 Specific activity-temperature relationship depends on substrate

The specific activities of both WT and two mutants (E316G and E361G) on substrate p-nitrophenol butyrate (C4) are shown in Figure 2a. As can be seen, WT has an optimal specific activity with C4 at 323K. The optimal temperature for E316G and E361G, mutants were 10 degrees higher at 333K. The specific activity data at temperatures below the optimum show that the two designed mutations increased the cold activity compared to the WT. Both WT and mutants have similar specific activities with C4 at T < 313K. At temperatures above their optimal temperature, the mutants have higher specific activities with C4 at most of the studied temperatures compared to WT. The E361G mutation increased the specific activity with C4 to a greater extent than the E316G. Moreover, mutant E361G had less change in specific activity (compared to E316G and WT) at T between 323K and 353K, indicating its relative higher tolerance to high temperatures and diminished temperature dependence.

**Figure 2.**
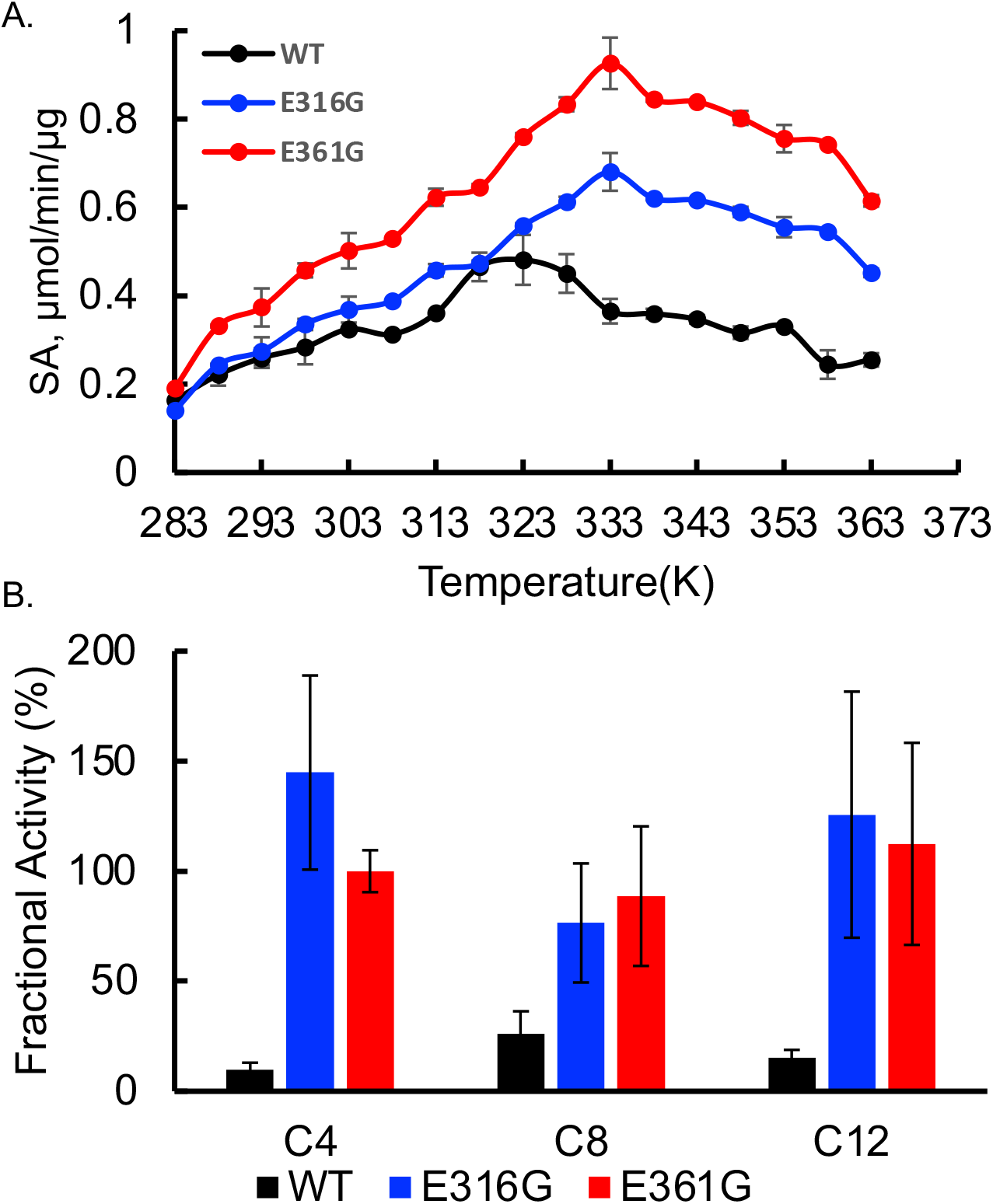
(a) Specific activity of WT (black), E316G (blue) and E361G (red) with C4 from T = 283 K to 363 K. (b) Kinetic stability of WT and the two glycine mutants measured as the % fraction of specific activity retained with C4, C8 and C12 at 323 K after 3 hours of incubation at 343 K. Error bars are standard deviation of at least three replicates.

The diminished temperature dependence of specific activity and shift in the optimal temperature correlate with the increase in the kinetic stability of the mutants (Figure 2b). This increase in specific activity and kinetic stability of mutants with C4 relative to WT suggests that flexibility inducing mutations can be introduced without compromising the thermophilic enzyme properties. In fact, in this particular instance, the mutations enhanced thermostability and thermophilic properties.

It has been shown that mutation around the catalytic triad in *Bacillus stearothermophilus* L1 lipase (BSL1) leads to changes in substrate specificity [29]. This change was attributed to the altered conformation and catalytic pocket [29]. As can be seen in Figure 3a, WT exhibited the highest activity at 313K with substrate p-nitrophenol octonoante (C8), which is different than its optimum temperature of 323K with substrate C4. The specific activity of WT showed a slow decrease from 313K to 338K, which was followed by an abrupt decrease when temperature reached to 343K. Compared to WT, both E316G and E361G had lower specific activity for most of the temperatures. E316G has higher activity than WT at temperatures below 293K. The optimum temperature for both E316G and E361G is 323K, which is 10K lower than the optimum temperature when substrate is C4, yet still 10K higher than WT. With substrate p-nitrophenol laurate (C12), both WT and E361G tend to have higher specific activities than with substrate C8 (Figure 3b). WT still have higher specific activity compared to mutants. Using C12, the optimum temperature of the WT was 313K, the E316G was 328K, and E361G was 333K.

**Figure 3.**
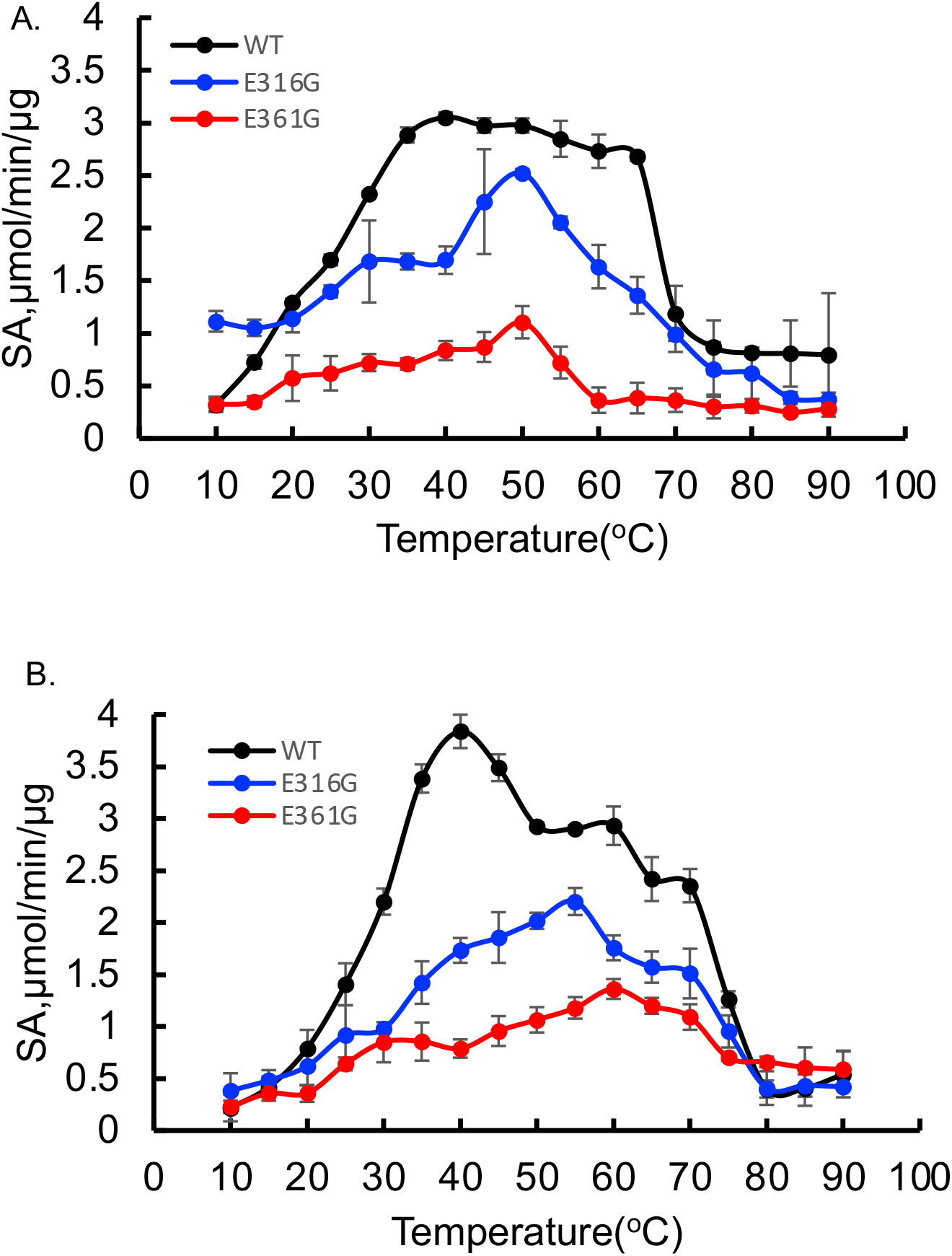
Specific activity of WT (black), E316G (blue) and E361G (red) with (a) C8 and (b) C12 at different temperatures. Error bars are standard deviation of at least three replicates.

The relative kinetic stability comparison between WT and mutants on C8 and C12 showed the same trend with on C4 where WT decreased significantly after 3 hours-thermal incubation at 343K while mutants still retain at least 80% of the activities (Figure 2b). Interestingly, analysis of the CD spectra of WT and the two mutants indicates similar overall secondary structure (Figure 4), suggesting the mutations effects on thermostability and specific activity are not caused by major conformational change. The CD spectra deconvolution indicated a 5% increase in alpha-helix, and 7% decrease in beta-strand and 2.9% increase in loop (turn and irregular) in the two mutants compared to the WT. (Supporting Information Table 3).

**Figure 4.**
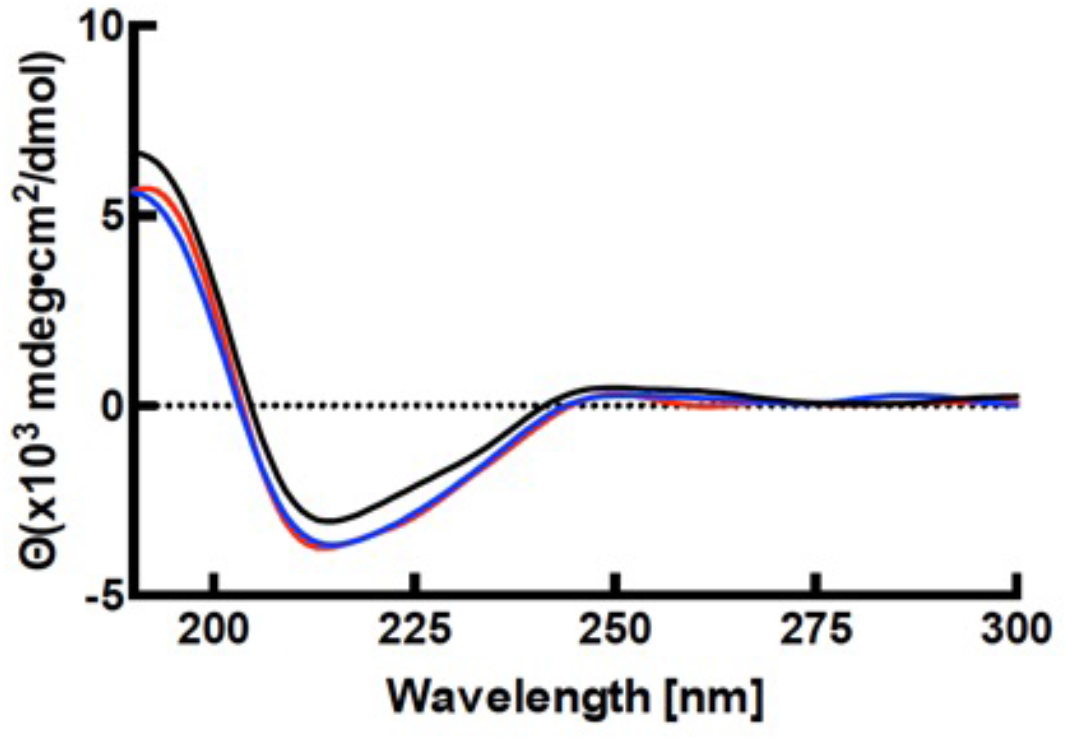
CD spectroscopy of WT (black), E316G (blue), and E361G (red) at T = 298 K. Purified enzymes were dissolved in 50 mM sodium phosphate at pH 7.0. The scanning wavelength was varied from 190 nm to 300 nm with 2 nm decrement. No significant changes in secondary structures of mutants were observed.

### 3.3 Psychrophilic traits in the mutants

The change in thermodynamic activation parameters Δ*H*^‡^ and Δ*S*^‡^ of mutants compared to WT obtained with C4 was determined using Eyring plots (Supporting Information Figure 1), which indicated both E316G and E361G have lower Δ*H*^‡^ and more negative Δ*S*^‡^ (Table 2) when compared to WT. These are typical characteristics of psychrophilic enzymes compared to orthologous mesophilic and thermophilic enzymes [2, 12, 18, 19]. Similar analysis using C8 and C12 revealed the same pattern of lower Δ*H*^‡^ and more negative Δ*S*^‡^ (Table 2) when compared to WT, irrespective of substrate and despite the differences in substrate specificity.

**Table 2.**
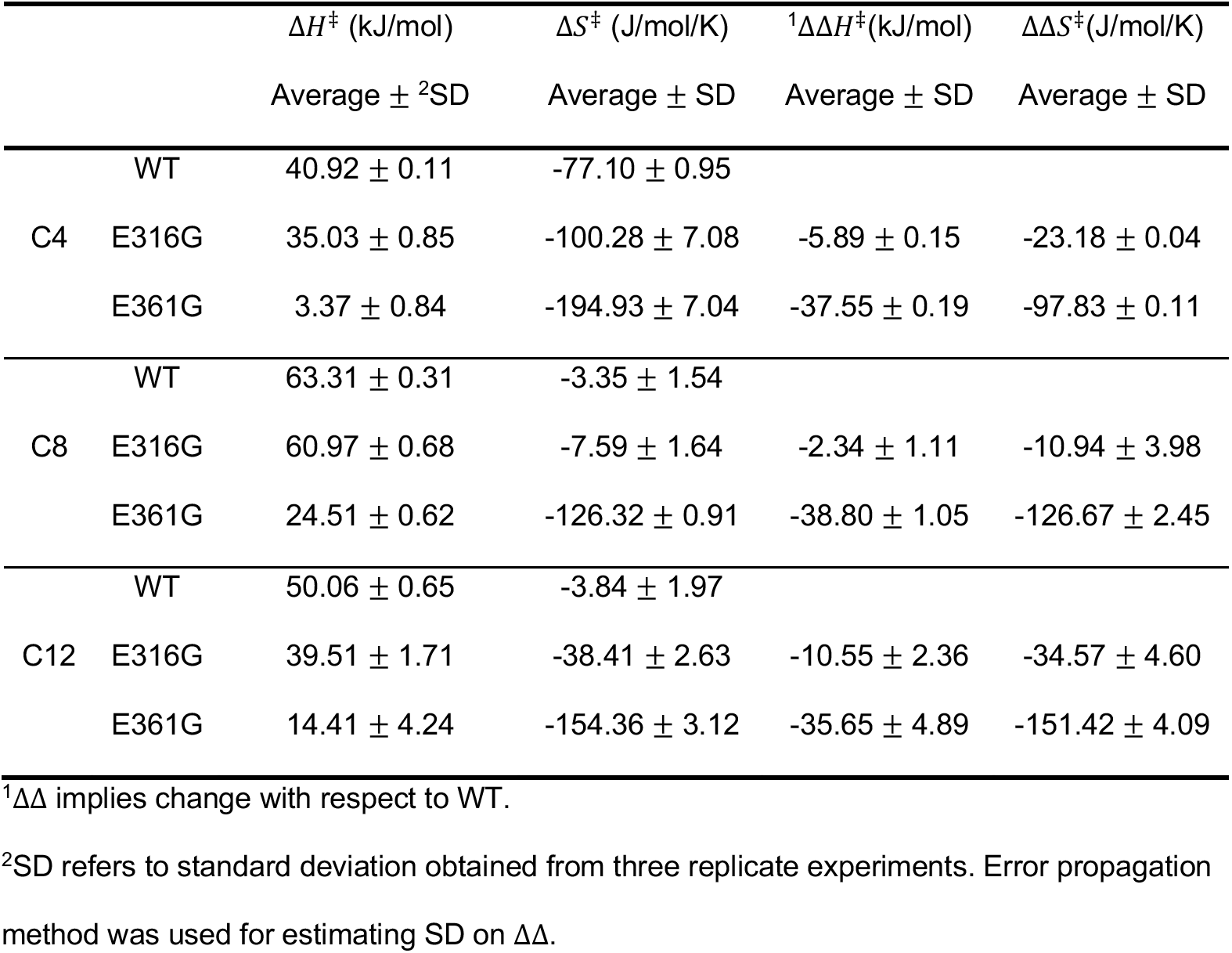
Δ*H*^‡^ and Δ*S*^‡^ of WT and mutants with C4.

The data show that E361G had much more different activation parameters compared to E316G and WT. In all substrates, the activation enthalpy was much smaller, and the activation entropy was far more negative. The E316G mutation had a smaller activation entropy and less negative activation entropy compared to the WT, but to a far lesser extent compared to the E361G mutant.

A comparison of the change in activation enthalpy and entropy for a given mutant as the substrate length is increased is more complex. For the WT and the two mutants, the activation enthalpy increases from C4 to C8 and then decreases from C8 to C12, where C4 < C12 (i.e., for Δ*H*^‡^ of a single enzyme, C4 < C12 < C8). A similar analysis of the activation enthalpy reveals a similar relationship, where Δ*S*^‡^ gets less negative from C4 to C8 and then gets more negative from C8 to C12 (i.e., for Δ*S*^‡^ of a single enzyme, C4 < C12 < C8).

For WT with C8 and C12 substrates, the Δ*S*^‡^ of WT is a small negative value (~0.003 kJ/mol/K), which suggests more similarity of ground state and transition state entropy of the enzyme-substrate complex in WT. Therefore, the Δ*G*^‡^ is largly dependent on the enthalpy changes, and consequently a higher exponential dependence on the catalytic rate with temperature according to Eyring, Evans and Polyani equation. This can explain the steep increase in the specific activity profiles of C8 and C12 (Figure 4) in WT with temperature. This is also observed on E316G with C8, where the Δ*H*^‡^ and Δ*S*^‡^ values, and specific activity profile closely follow that of WT.

Deviations in Δ*H*^‡^ and Δ*S*^‡^ are larger between E316G and WT with C12, where the correlation in the differences are higher with that of psychrophilic and thermophilic enzymes. The lesser increase in specific activity with temperature in E361G with C8 and C12 is consistent with this reasoning where decrease in Δ*H*^‡^ and increase Δ*S*^‡^ essentially decreases exponential dependence of catalytic rate with temperature. Nevertheless, the differences in Δ*H*^‡^-Δ*S*^‡^ primarily explain exponential temperature dependence of activity with temperature but not the dependence of specific activity on the substrate size. Further, the changes in the trends of Δ*H*^‡^-Δ*S*^‡^ with substrate size indicates an evident role in the ability of the enzyme to accommodate the substrate within the binding site.

## 4.0 Discussion

Our central hypothesis was that by introducing flexibility inducing mutations near the active site we could introduce cold activity by decreasing the temperature dependence of *k_cat_*. Our data demonstrated two mutations, E316G and E361G, could enhance cold activity in GTL; however, these mutations led to a few unexpected changes, such as increased thermostability and altered substrate specificity. These data support the overall hypothesis that cold activity can be rationally designed by mutations that lower the activation enthalpy and make the activation entropy more negative.

There are several theories attempting to explain the molecular mechanisms underlying cold activity. Flexibility around active sites has been considered responsible for the cold activity adaptation of psychrophilic enzymes. Feller et al. proposed that psychrophilic enzymes need to increase flexibility of the active sites and maintain stability of other domains in order to reduce the counter effect of the more negative activation entropy Δ*S*^‡^ [17, 19]. A recent protein softness theory suggests temperature adaptation of enzymes is believed to be mediated by protein surface softness [4, 13, 23, 30–32]. Activation enthalpy and entropy values gradually became thermophilic when the restraint moves deeper from the protein surface towards the internal core[^4, 23]^. However, a temperature adaptation study of *Bacillus agaradherans* cellulase Cel5A revealed fluctuation on active sites could be enhanced and further increase the robust activities at low and warmer temperatures by disrupting the hydrogen bonds around the catalytic residues [27]. Together this suggests that the psychrophilic/thermophilic traits transition might not be simply regulated by local flexibility (surface softness or active site fluctuation). Instead, they are regulated by the allosteric residues which tune the overall structure fluctuation and further the entropy/enthalpy balance. As has been proposed by Hacisuleyman et al., surface residues tend to become entropy sources and drive the fluctuations of other residues inside the protein core [33]. Agarwal et al. proposed that charged residues with long chain on the protein surface tend to form interactions with solvent and regulate catalysis [34, 35]. This has been corroborated by the study on adenylate kinase in which its cold adaptation was driven by dynamic allostery on the surface domains [36], leading to propagation of the fluctuations from the mutated residues to the overall structural change. This also explains why the secondary structure of WT and mutants are similar while their temperature adaptations are significantly different.

To explain the higher relative specific activities of E316G and E361G on substrate C4, kinetics assay was employed to investigate whether the activation enthalpy and entropy have transformed into psychrophilic-like traits. As shown in Table 2, both mutants showed significant decrease in activation enthalpy and more negative entropy values. These changes indicate the inclusion of psychrophilic traits in mutants.

The impact of the mutations on the substrate specificity were possibly due to the influence of the substrate hydrophobicity on the lid opening. Lipases are typically activated at hydrophilic-hydrophobic interfaces, which are more likely to be induced by the longer substrates [37]. The relatively lower activity of GTL mutants on longer substrates might also be explained by either reduced lid opening. The perceived enhancement of activity on C4 substrates is more properly classified as less decrease in activity, as the specific activity values were at least 4-fold lower with C4 compared to C8 or C12. If the GTL mutations effectively diminished the effect of hydrophobic interfaces in lid opening, this would be consistent with the experimental data. Indeed, Dror et al., [38] observed that reducing the polarity of residues on the lid had exactly this effect. The mutations of acidic residues to glycine in proximity to active site, where the lid interacts in closed form, would likewise reduce the polarity of lid interactions.

The unexpected increase in thermostability is difficult explain from a structural perspective without further studies. In a study of metagenome derived lipase, the paralogous mutation of E315G (similar to E316G in GTL) resulted in reduced thermostability [29]. One hypothesis supporting the increased thermostability of E316G GTL (and E361G), is that since the enhanced flexibility is located outside the active site, it can be effectively dissipated through allosteric interactions that ultimately rigidify other regions of the enzyme and effectively ratchet the lipase into a more thermostable configuration, perhaps by forcing the lid open. Furthermore, the Dror study [38] on reducing the polarity of the lid also noted increased thermostability, which would be consistent with our observations.

To summarize, our kinetics assay has investigated engineering active site flexibility as a means of altering the temperature dependence of catalysis and increasing cold activity in an otherwise thermostable enzyme. The activation enthalpy and entropy measured provided proof that the rationally designed mutations resulted in psychrophilic adaptations compared to the WT. Previous studies of enzyme temperature adaptation tend to relate the enzymatic catalysis over temperature with the local structural change [18, 26]. Our work found temperature adaptation of enzymes that result in modest structural change could have a profound impact on optimal temperature shift, enhanced thermostability, and altered substrate specificity. Proof that the impact of these simple rational mutations induced active site proximal flexibility still needs verification from molecular dynamics simulation. Nevertheless, we successfully engineered psychrophilic traits into a thermophilic enzyme through a single mutation around the active sites without sacrificing its thermostability. Our work provides an alternative approach for enzyme temperature adaptation engineering.

## 5.0 Author Contributions

**Weigao Wang**: Conceptualization, Methodology, Investigation, Writing – Original Draft, Writing – Review & Editing. **Siva Dasetty**: Conceptualization, Writing – Original Draft, Writing – Review & Editing. **Sapna Sarupria**: Conceptualization, Writing – Original Draft, Writing – Review & Editing, Supervision. **Mark Blenner**: Conceptualization, Methodology, Writing – Original Draft, Writing – Review & Editing, Supervision, Project Administration, Funding Acquisition.

## 6.0 Acknowledgements

The authors thank Max Hilbert for collecting circular dichroism data. The authors thank the following individuals for helpful comments and edits: Molly Wintenberg, Dyllan Rives and Allison Yaguchi. The authors thank Bob Latour for access to the circular dichroism polarimeter, which is supported by SCBioCRAFT (NIH-2P20GM103444). The authors acknowledge funding from the Air Force Office of Scientific Research to MAB (FA-9550-15-1-0163).

---

**Supporting Information Table 1.**
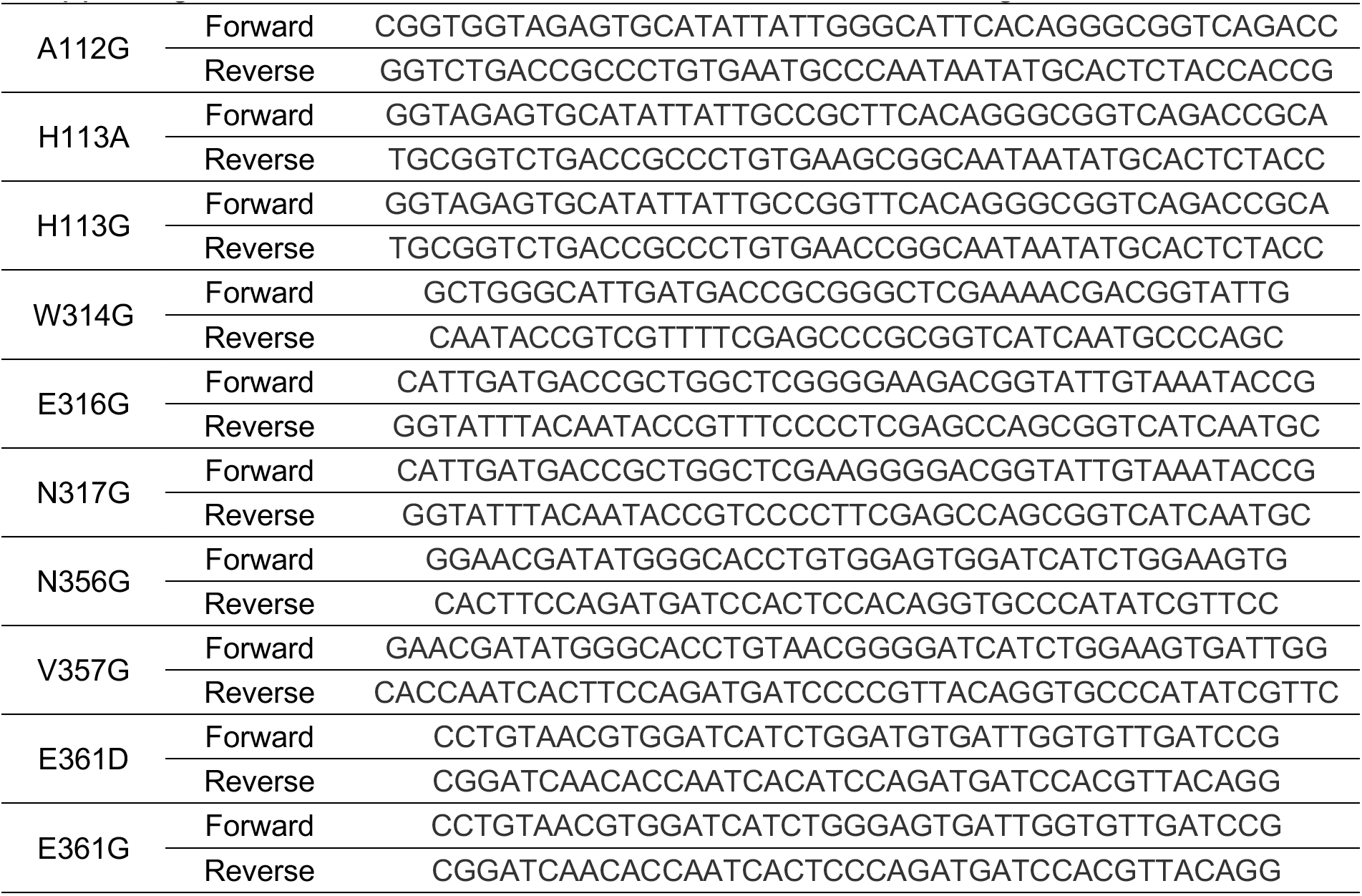
Primers for site directed mutagenesis

**Supporting Information Table 2.**
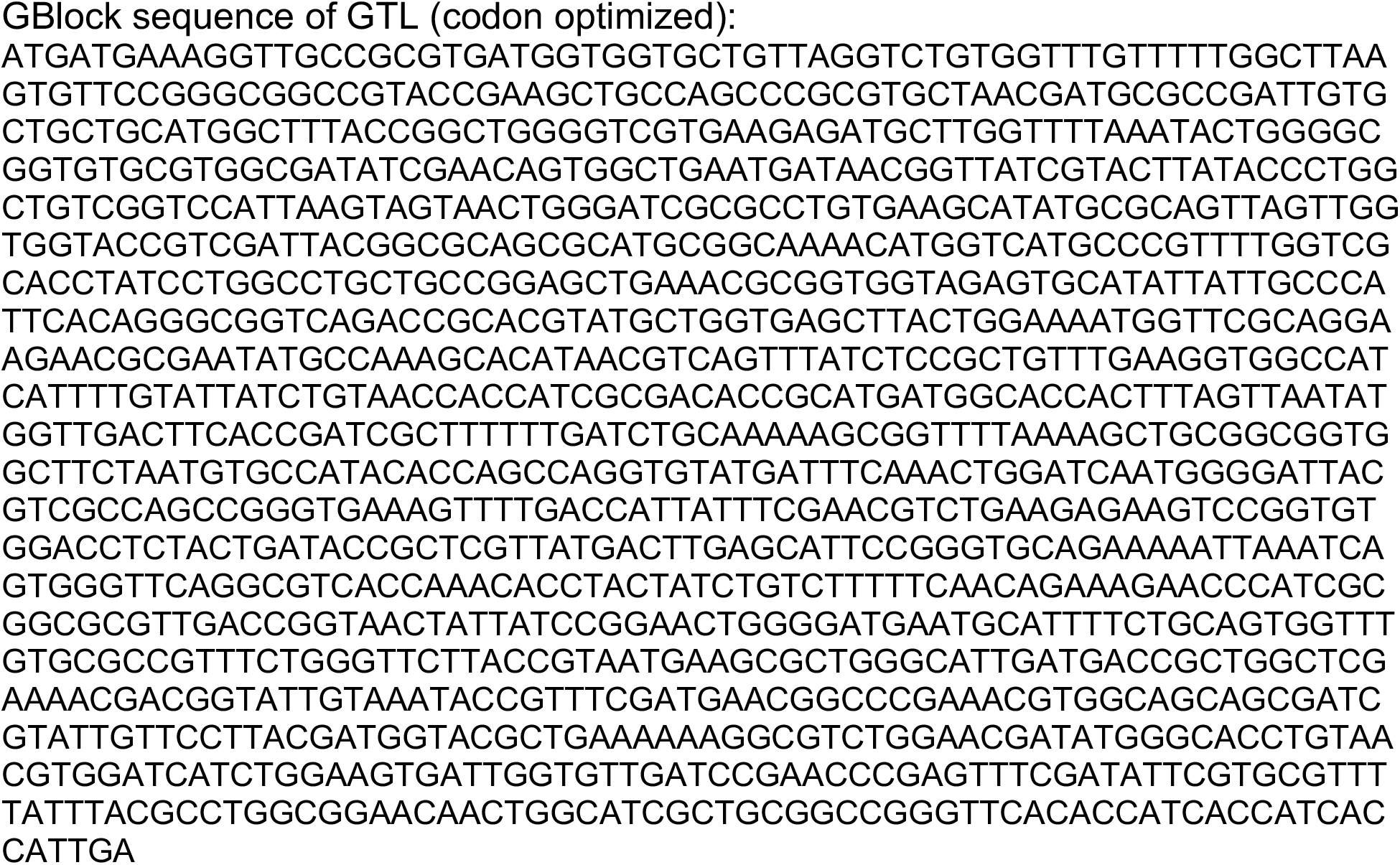
Primers for site directed mutagenesis

**Supporting Table 3.** Deconvolution of Circular Dichroism Data.

|  | WT | E316G | E361G |
| --- | --- | --- | --- |
| Helix | 0.285 | 0.236 | 0.236 |
| Beta Strand | 0.221 | 0.297 | 0.297 |
| Turn | 0.195 | 0.195 | 0.195 |
| Irregular | 0.271 | 0.277 | 0.277 |

**Supporting Figure 1.**
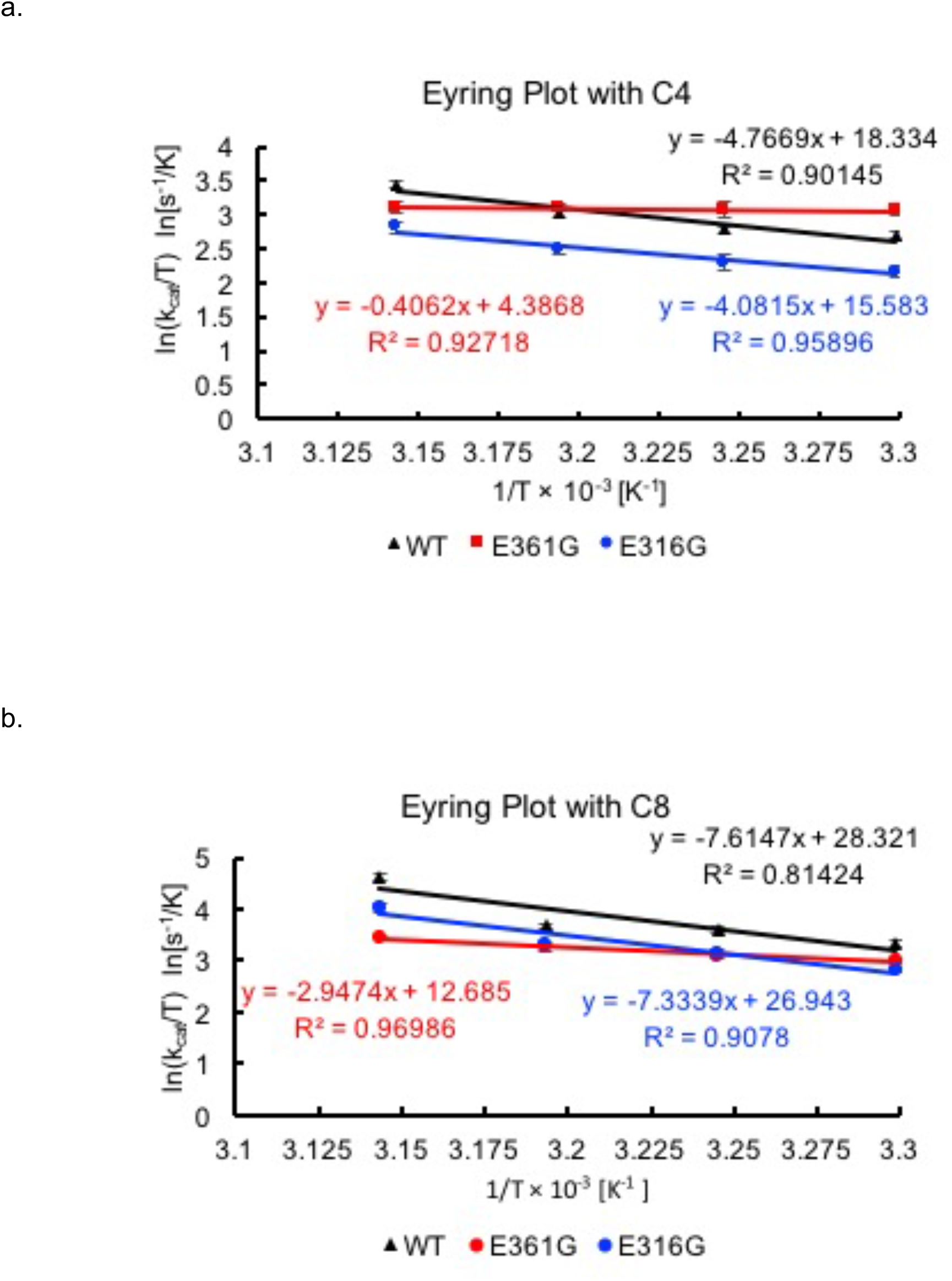
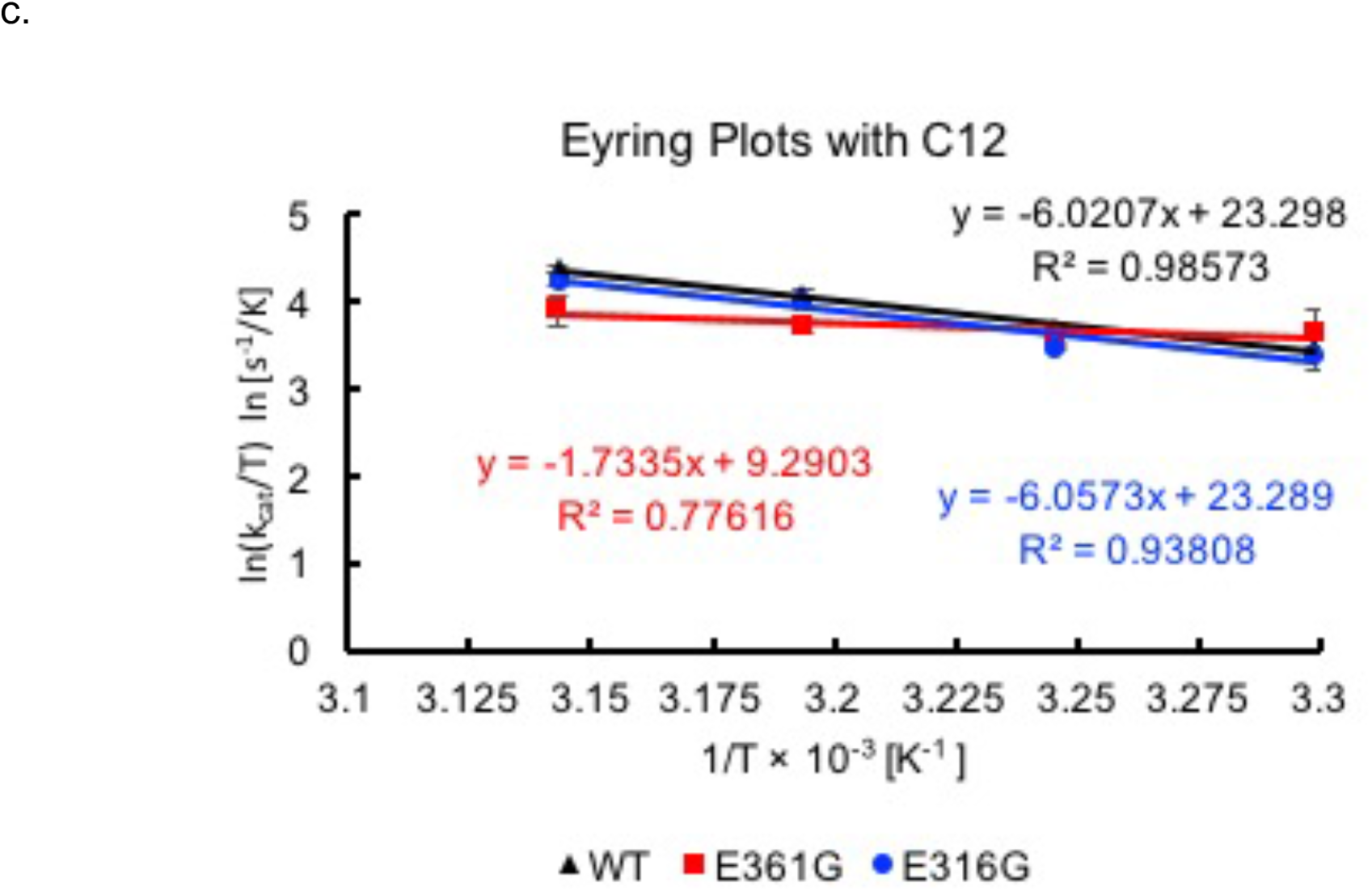
Eyring Plots Used to Calculate Thermodynamic Parameters

